# Sustained Selective Attention in Adolescence: Cognitive Development and Predictors of Distractibility at School

**DOI:** 10.1101/2023.01.15.523576

**Authors:** Michael H. Hobbiss, Nilli Lavie

**Affiliations:** Institute of Cognitive Neuroscience, University College London

**Keywords:** selective attention, sustained attention, response variability, adolescence, school education

## Abstract

Despite much research into the development of attention in adolescence, mixed results and between-task differences have precluded clear conclusions regarding the relative early- or late-maturation of attention abilities. Moreover, although adolescents constantly face the need to pay attention to their lessons at school, it remains unclear whether laboratory measures of attention can predict their ability to sustain attention focus during school lessons. Here we therefore devised a task that was sensitive to measure both sustained and selective attention and tested whether any of our task measures can predict adolescents’ levels of inattention during their school lessons. 166 adolescents (aged 12-17) and 50 adults performed in our sustained-selective attention task, searching for letter targets, while ignoring salient yet-entirely-irrelevant distractor faces, under different levels of perceptual load-an established determinant of attention in adults. Inattention levels during a just-preceding classroom lesson were measured using our novel self-report classroom-distractibility checklist. The results established that sustained attention (measured with response variability) continued to develop throughout adolescence, across perceptual load levels. In contrast there was an earlier maturation of the effect of perceptual load on selective attention: load modulation of distractor interference was larger in the early adolescence compared to later periods. Both distractor interference and response variability were significant unique predictors of distractibility in the classroom, including when interest in the lesson and cognitive aptitude were controlled for. Overall, the results demonstrate divergence of development of sustained and selective attention in adolescence, and establish both as significant predictors of attention in the important educational setting of school lessons.

## Sustained Selective Attention in Adolescence: Cognitive Development and Predictors of Distractibility at School

The ability to focus attention on goal-relevant information whilst ignoring irrelevant distractions, and maintain this focus throughout a mental task, is critical for effective information processing across perception, learning and memory. Children in particular face the challenge of keeping their attention focused at class, or home study sessions, throughout their schooling years.

Understanding attention development, together with establishing whether experimental laboratory measures can apply to predict levels of attention focus during lessons is therefore highly valuable for a comprehensive developmental theory and its potential applications. However, while much is known about the development of attention in early childhood, the maturation process of attention in the adolescence period remains somewhat puzzling: some studies have suggested attention has maturated already by early adolescence, while other studies have demonstrated continued development until adulthood. It is plausible that different attention abilities develop at different rates (for example due to the extent to which they each rely on frontal cortex regions that continue to mature throughout adolescence). However, it is important to establish such a developmental divergence within the same task, otherwise task-differences can potentially account for the different developmental results. Nevertheless, previous research has typically used different paradigms to study either ‘selective attention’ or ‘sustained attention’ and together with the great variety of stimuli used in the different studies a clear resolution remains elusive.

In the present study we therefore devised a ‘sustained selective attention’ paradigm, which allowed us to investigate both selective and sustained attention abilities within the same task, and test whether our task measures of both attention abilities can predict adolescents’ attention in the school classroom. Further by relating our research to important determinants of attention in adults, we hypothesized that the nature of the distractor stimuli and the level of perceptual load in the task might be critical factors, that can allow us to resolve previous discrepancies.

Next, we briefly review the relevant previous research, highlighting the inconsistencies and how these relate to potential determinants of attention in adolescence.

## Evidence for Mature Selective Attention in Adolescence

Adolescents’ development of selective attention, has typically been studied with the ‘flanker task’ (Eriksen & Eriksen, 1974) in which a target letter, or shape (e.g. an arrow) is presented in the display centre, flanked by one or more distractors that are either associated with the same (in ‘congruent’ conditions) or the other target response (incongruent conditions).^1^ The findings typically revealed no developmental effects. For example, Grose-Fifer et al. (2013) found no difference in distractor congruency effects in a letter-based flanker task, between adolescents (aged 15-17) and adults. Huizinga, et al. (2006) used an arrow-based flanker task and similarly found that while both younger (aged 6-8) and older children (aged 10-12) displayed larger congruency effects on RT than mid-adolescents (aged 14-16), and adults (aged 18-26), congruency effects did not differ between mid-adolescents and adults. An ERP study using the same arrow-based flanker paradigm also showed no behavioural difference in congruency effects between adolescents (aged 16) and adults, with both groups responding more quickly and accurately irrespective of the distractor congruency conditions than a group of 12 years old (Ladouceur et al., 2007). Gyurkovics, et al. (2020) also used an arrow-based flanker task, and found that both the flanker congruency and congruency-sequence effects were unaffected by age, across groups of 12–13, 14–15, 18–20 and 25–27 years old.

In an exception, a flanker task in which choice responses to a central target circle were made based on the its colour) adolescents up to age 14-15 showed larger congruency effects than adults (Waszak, et al., 2010). These results may have been due to the higher visual salience of the colour match/mismatch, especially given that four flanking circles-surrounded the target circle (all either matching or mismatching with its colour). Thus for attention tasks with distractor stimuli (such as letters or arrows) that are not particularly salient the ability to ignore a response-competing distractor, which is known to develop in childhood (e.g., Huang-Pollock et al., 2002), appears to reach maturity by adolescence. In contrast, flanker tasks involving affective, or motivationally-relevant distractors such as happy vs. fearful congruent or incongruent distractor face expressions greater congruency effects are found in both young (11-14) and older adolescents (15-17) compared to adults (Grose-Fifer et al. 2013).

Similarly in another measure of selective attention based on the phenomenon of attentional capture by the presence of a unique ‘singleton’ distractor (e.g. a red among all green shapes, in a shape-based visual search task) adolescents attentional capture was not found to be larger for adolescents (aged 13-16) in comparison to adults, as long as the singleton distractor was not previously associated with a reward (through a ‘value learning’ procedure, see Anderson et al., 2011). Only the reward-associated singleton distractors were found to produce stronger and longer lasting attentional capture effects on visual search in adolescents compared to adults (Roper et al., 2014).

One explanation for the findings that typical selective attention paradigms only reveal larger impact of distractor when these are either affective or motivationally salient may be in terms of reduced emotional regulation in adolescence (Heller & Casey, 2016; Larson et al., 2002), or reduced cognitive control over task priorities (i.e. prioritising motivationally-relevant previously rewarding stimuli over the neutral task-target stimuli), rather than an immature selective attention ability per se (see Lavie, 2005; Lavie et al., 2004, for further discussion of this distinction). Alternatively, it is possible that selective attention is not fully developed in adolescence, but that typical flanker or attentional capture tasks are less sensitive to reveal this, because their distractor stimuli are typically not sufficiently salient (c.f. Waszak, et al., 2010), or meaningful (e.g. an arrow conveys a rather restricted meaning compared to an emotional face, or a stimulus associated with reward) to reveal any reduced ability to focus attention. We therefore concluded that a sensitive measure of selective attention development should involve irrelevant distractors that are salient and meaningful but emotionally and motivationally neutral.

## Perceptual Load

Selective attention is known to critically depend on the level of perceptual load in the current task (Lavie, 1995; Lavie & Tsal, 1994). High perceptual load in a task (e.g. tasks involving a larger number of stimuli, or increased demand on perception for the same number of stimuli, e.g. conjunction vs. feature search) has been shown to allow a higher level of attentional engagement, increasing neural signal associated with task-relevant stimuli and reducing the neural response to task-irrelevant distractors (e.g. Bruckmaier et al., 2020; Torralbo et al., 2016). These effects are explained within the load theory account for attention, in which attention effects are explained with the combination of a limited capacity resource approach with automatic allocation of all the resource on all stimuli within the limited capacity. Cognitive control is restricted to maintaining processing priorities (so that higher priority is given to task-relevant stimuli), however, if the task processing does not take up full capacity, spare resources are allocated in parallel, involuntary manner to the irrelevant stimuli that people wish to ignore, resulting in distractor interference effects. Perceptual load in the task processing is therefore a critical determinant of selective attention. High perceptual load that exhausts capacity in task-relevant processing, results in the selective processing of task relevant information only, with irrelevant distractors effectively filtered out. However, in task conditions of low perceptual load attention will automatically spill over to the processing of irrelevant distractors as well. The effects of perceptual load have been established across different perceptual load manipulations and measures of distractor processing (for reviews see Lavie, 2005; Murphy et al., 2016). The effects of perceptual load on selective attention have been shown to apply across individual differences in the level of distractibility (e.g. Forster & Lavie, 2016) and perceptual load has also been shown to improve selective attention focus in ADHD (Forster et al., 2014) and in typically developing children (Huang-Pollock et al., 2002), while both populations display greater susceptibility to distraction in comparison to healthy adults. Given that when the distractor stimuli are affective or motivationally-relevant adolescents have also been found to have elevated levels of distractibility (compared to adults), and may also be more susceptible to distractions from salient, and meaningful yet irrelevant distractors, we sought to examine whether adolescents can also benefit from increased perceptual load to improve their ability to focus attention in the presence of such distractors.

## Development of Sustained Attention in Adolescence

Research of sustained attention has typically used different tasks to those used to measure selective attention such as the Continuous Performance Test and the Sustained Attention to Response Task (SART). Both tasks involve monitoring a sequence of stimuli (e.g. digits) appearing one at a time, and responding to some of them while withholding responses to other stimuli (nontargets in the CPT and ‘no-go’ targets (e.g., the digit 3) in the SART. The results indicate that in addition to increased rate of failures to inhibit responses (or commission errors), adolescents (up to age 16 in some of these reports) also display elevated RT variability in comparison to adults (Braet et al., 2009; Fortenbaugh et al., 2015; Gyurkovics et al., 2020; Stawarczyk et al., 2014). While the inhibition failures may be attributed to their immature inhibitory control, the increased response variability in their routine go responses, can provide a purer measure of adolescents’ ability to pay attention irrespective of an immature inhibitory control. Increased variability has also been associated with greater rates of mindwandering (Cheyne et al., 2009; Stawarczyk et al., 2014) suggesting that both are driven by a failure to sustain attention focus on the task. Indeed Cheyne et al. (2009) have demonstrated that increased variability in the SART is not only associated with increased mindwandering reports, but also with increased self-reported attentional failures.

It remains unclear, however, whether adolescents will show reduced sustained attention ability when completing a focused attention task under higher perceptual load that is known to be more attention engaging. Specifically, both the CPT or SART tasks involve a low level of perceptual load (monitoring for clearly distinct targets, each appearing singly on the screen, and with fairly slow presentation rates), while in our task conditions of high perceptual load participants perform a visual search task with multiple items, searching for a target among similar non-target stimuli. The wealth of studies showing that people are able to fully focus attention on the task in conditions of high perceptual load (e.g. Lavie, 1995, Lavie & Cox, 1997; Lavie & Fox, 2000, Lavie, 2005 for review), and that high perceptual load improves focused attention in ADHD (Forster et al., 2014) and in childhood (Huang-Pollock et al., 2002) raise the important question of whether adolescents would also be capable of sustaining attention focus in tasks condition of high perceptual load at a younger age than that reported based on their performance in the CPT or SART tasks of low perceptual load.

## Predicting Adolescents Distractibility in the Classroom

As we mention earlier understanding adolescents’ ability to sustain a selective focus of attention is of paramount importance also given the critical role attention plays in learning and education, and the significant educational milestones adolescents face in their secondary school period. Indeed, a good deal of educational research has linked teachers’ ratings of inattention in the classroom to educational achievement (e.g., Gray et al., 2017). However teacher ratings of inattention can vary in their reliability both within and between schools (Merrell & Tymms, 2001). For example, some studies reveal that fewer than half of the pupils who receive clinically elevated ratings of attention problems from their teachers in a given year also maintained this level of rating in the following year (Rabiner et al., 2010). This points to the need to augment teachers’ ratings with task-based measures of attention.

Some previous research has related attention task performance to teachers’ ratings of inattention in the classroom, however so far this research has only been conducted with children. For example, Johnson et al., (2020) found that 8 and 11 years old children’s commission errors in the SART were positively correlated with teacher ratings of both inattention and hyperactivity-impulsivity, and negatively correlated with on-task behaviour in the classroom. A measure of response variability was also negatively associated with on-task classroom behaviour. Other measures of inhibitory control (such as the Stroop, and visual and auditory response competition tasks, where participants respond to target stimuli whilst inhibiting responses to other similar stimuli) have been found to correlate with teacher ratings of pupils’ attention control skills (at least in terms of differentiating between the top and bottom quartiles) in children (aged 8-12), (Das et al., 1992; Papadopoulos, et al., 2002). A negative correlation between CPT performance and the inattention subscale of the Conners Teacher Rating Scale was also found for 7 and 10 years old children (Lam & Beale, 1991). These findings therefore provide preliminary evidence of an association between cognitive measures of attention control and some measures of classroom inattention; however all these studies have concerned the childhood period. Thus while prior work is promising regarding the use of task measures to predict distractibility in the classroom, it remains unclear whether any immature attention abilities that remain in adolescents can be predictive of their ability to focus attention in the classroom.

## The Present Study Aims and Task Paradigm

In the present study we thus set out to investigate: i) whether any developmental trajectories of the abilities to both focus selective attention, and to sustain a stable focus, remain in adolescence when these are measured in an attention task that is sensitive to depict both attentional functions. ii) whether our attention task measures can be used to predict the level of focused attention in the real-life setting of a lesson in the school classroom. iii) whether increased perceptual load in the task would modify any of the developmental trajectories or their predictive value.

A promising task-based measure of the type of inattention that may harm learning is the irrelevant distraction paradigm (Forster & Lavie, 2008A). This task paradigm has been designed to measure the interference caused by distractors that are salient and meaningful, yet entirely irrelevant to the task, in order to provide a measure of selective attention that can better reflect (in comparison to the prevalent selective attention paradigms of response competition and attentional capture) the mechanism of focused attention, and conversely distractibility, as it functions in the real world.

In both previous paradigms the distractors are not task-irrelevant, but instead are either directly associated with the task response, in the response competition paradigm (e.g. congruent or incongruent), or include some search-relevant features, in the attentional capture paradigm (for example, both the search target and the distractor involve a ‘singleton’ feature, e.g. being the only odd shape, or odd colour, in an otherwise homogenous search array). In the real world, distractors are often entirely irrelevant to the task, yet they can disrupt attention focus and thus interfere task performance. For example, a pupil might be distracted by the sight of another pupil walking by the classroom window, and make errors in their current math exercise task.

Forster and Lavie reasoned that the task interference effects from such distractors occurs, despite their task-irrelevance due to a combination of both bottom up (e.g. greater visual salience relative to the task stimuli) and top down factors (e.g. being more meaningful, or less expected than the task stimuli, see Forster & Lavie, 2008B) which makes the distractors strong competitors for attention even though they are not directly competing with either the task response or the search target (c.f. the response competition or attentional capture paradigms). Therefore, an attention task that combines these factors in its distraction measure should provide a sensitive measure of selective attention, that is capable of predicting inattention failures in real life situations, and as experienced in individuals that have a higher propensity to being distractible (e.g. clinically diagnosed with ADHD, see Forster et al., 2014; or along the spectrum of individual differences in inattention in non-clinical populations, see Forster & Lavie, 2016).

The irrelevant distractor paradigm was therefore designed to include all these elements as follows. Participants perform a visual attention task (e.g. letter-based visual search task, or a continuous choice response task discriminating between letters and digits, e.g. Forster & Lavie, 2008A,B; Forster & Lavie, 2011), while being instructed to ignore any irrelevant distractors. On a minority of the task trials (e.g. 10%), an entirely task-irrelevant distractor image that is both visually salient and meaningful (e.g. of a famous cartoon characters, or a human face), appears in a task-irrelevant location, e.g. in the display periphery, (see Forster & Lavie, 2021 for a recent demonstration). Distractor interference effects are measured on the basis of the cost to the attended task RT in the presence versus absence of this task-irrelevant distractor.

In addition to predicting the aforementioned individual propensity to inattention failures in daily life (Forster & Lavie, 2016), distractor interference effects in this task have also been shown to predict susceptibility to mindwandering (Forster & Lavie, 2014). Since inattention failures in the classroom are expected to reflect distractibility by task-irrelevant sources, triggered both by external task-unrelated stimuli (e.g. another pupil, as in the example outlined earlier), and internal sources such as mindwandering, as well as inherent fluctuations in the ability to sustain attention on task (e.g. see Esterman et al., 2014), we have used the irrelevant distractor task to measure both selective attention (based on the distractor interference effects) and sustained attention (based on reaction time variability in the absence of a distractor) in adolescents (compared to adults) within the same task paradigm, under different levels of perceptual load, and examined whether our task measures of selective and sustained attention, can predict classroom inattention.

In order to measure adolescents experience inattention in the classroom, we modified a self-report measure of everyday experiences of distractibility and mindwandering (Hobbiss et al., 2019) to be suitable to measure these in the classroom.

## Method

### Participants

50 (33 F) students, aged 20-35 (M=24;6, SD =4;3), volunteered through the UCL Psychology subject pool, and 166 secondary-school students participated: 59 (30 F) in the ‘early adolescence’ age group aged 12-13 (M=13;0, SD =0;6), 73 (28 F) in the ‘mid-adolescence group’ aged 14-15 (M=14;10, SD =0;6), and 34 (18 F) in the ‘later adolescence’ group aged 16-17 (M=17;3, SD =0;3 years).

Adolescent sample sizes were determined using a power analysis using G*power (Erdfelder et al., 2009), based on Forster et al. (2014), who found a significant 2×2 interaction between group and distractor condition (η^2^_p_ =.231). This effect size suggested that a total sample size of 132 (i.e., n=33 per group) would provide 95% power to detect between-group differences across four comparisons (3 comparisons between the 4 age groups, as well as a potential comparison of the whole adolescents group to the adults) with a multiple comparisons correction (alpha =0.0125). During the testing period, it became possible to exceed these sample size targets.

Participants scoring chance level accuracy (50%) or below in any experimental condition were excluded (n =9), as were those with reaction times more than 3 standard deviations from the mean (n =1). The exclusions were spread across the ages (13-14yrs =5, 14-15yrs =2, 16-17yrs =2, adults =1), leaving the final samples of 49 adults (20-35, M=24;0, SD =4;1) and 54 (28F) adolescents in the ‘early adolescence’ group (M=13;0, SD =0;6), 71 (27 F) in the ‘mid-adolescence’ group (M=14;10 months, SD=0;6), and 32 (17F) in the ‘later adolescence’ group (M=17; 3, SD=0;3).

Informed consent was obtained for all participants (parental consent for adolescents). The research was approved by UCL research-ethics committee.

### Stimuli and Procedure

Adult participants were tested on site at UCL. Adolescent participants were tested at school, immediately after a school lesson. In both sites testing took place in a small, darkened, single-subject room. The experiment was run using MATLAB (Mathworks, Inc.). Participants established 60 cm distance from their chin to the monitor, using a 60 cm long string attached to the monitor, and were asked to maintain this distance throughout the experiment.

Participants performed a visual-search task, searching for either ‘X’ or ‘N’ in a circle of six dark-grey letters presented at the display centre on a light-grey background. In low load displays the five non-target positions were occupied by lower-case ‘o’s. In high load displays they were occupied by five different angular letters from a list of ten (H, K, M, W, T, V, L, Z, F, E). Target position and identity were fully counterbalanced within blocks for each load condition. A face distractor subtending 2.3° vertically was presented at fixation on a random 10% of trials at a minimum of 1° edge-to-edge distance from the nearest letter. 18 different adult faces of a neutral-expression (and open mouth, since these have been shown to be closest to the neutral ratings for both females and male faces; e.g. Langeslag et al., 2018) were selected from the NimStim battery (Tottenham et al., 2009). Each face appeared only once in each of the load conditions. Participants were asked to place and keep their fingers on the 0 and 2 keys of the numeric keypad throughout each block, and press 0 for ‘X’ and 2 for ‘N’ as fast as possible, while remaining accurate. They were also asked to ignore the distractor faces, and this was emphatically stressed both in the instructions on the computer screen, and by the experimenter.

Participants completed six blocks of 60 trials each (3 per each load condition) preceded by 18 example trials (9 per each load). Participants alternated between and high and low load blocks in a counterbalanced order of ABBAAB or BAABBA. Half of the participants began with a low-load block (A), and half began with a high-load block (B).

Each block was preceded by three ‘warm-up’ trials, which never contained a distractor, and were excluded from the analysis. Each trial started with a 1000 ms central fixation cross, which was replaced by a 200 ms search display, followed by a 3000 ms blank response screen. A new trial was initiated following a participant’s response, or 3 seconds if they failed to respond. A ‘beep’ tone provided feedback for errors and missed responses.

Participants completed a self-report distraction checklist (either before or after the experiment, counterbalanced across participants) about their experience of distractibility in the classroom lesson immediately preceding the experiment^2^. The distraction checklist was adapted from a previous study (Hobbiss et al., 2019) to the school classroom, based on an eight-teachers focus group who discussed prominent sources of distraction in lessons. The teachers were asked initially to provide as many possible sources of classroom distraction having been present in their lessons. A second stage of the discussion then looked to create more general distraction categories from the specific distraction items discussed (For example, the items of ‘tapping’, ‘humming’ and ‘background talking’ were amalgamated into a category of ‘background noise’). Items which did not clearly identify an external source of distraction were discarded, these included ‘tiredness’, ‘boredom’ and ‘cognitive overload’. All the categories selected for the checklist were based on the majority of the group agreement that the category “could present a significant source of distraction for students in average lessons”.

This produced seven categories of distraction sources: ‘people around you (friends, classmates, teachers etc)’, ‘background noise (music, talking, tapping, shouting etc)’, ‘Social networking (alerts, posts, updates, messages etc)’, ‘other functions/apps on your phone/tablet/computer (games, internet browsing etc)’, ‘displays in the classroom (posters, maps, artwork etc)’, ‘unrelated thoughts (mindwandering)’ and ‘looking out of the window’. For each of the different distraction sources, participants reported for how long their attention was diverted away from their main focus towards the distractor, using a slider bar from 0-15 minutes, and text box on its right for participants to write a time if more than 15 minutes. The range of responses was 0-50 (the maximum lesson duration). Participants were instructed that it was possible to report multiple distractions occurring simultaneously (so that the total duration reported did not need to add up to 50 minutes). The self-report checklist also included questions about participant’s age, gender, interest in the topic being studied, the subject, and whether the lesson involved working alone (or with others) or work on a computer.

### Cognitive Abilities

Results of the adolescents’ performance on the Cognitive Abilities Test (CAT 4^th^ Edition, GL Assessment, 2016) were supplied by the school. The CAT is a widely used test of intelligence and reasoning skills in the UK and Ireland, consisting of four batteries of multiple-choice questions assessing verbal, non-verbal, quantitative and spatial reasoning, which can be combined into an ‘global’ cognitive ability score. It was this global score that was supplied by the school.

### Statistical Analyses

#### Experiment measures of Attention

Our main experiment measure of selective attention were based on mean RT (as is customary in this paradigm) and the null hypothesis significance testing (NHST) results. Bayesian statistics are reported as well for any key effects found as non-significant in the NHST, in order to assess the level of evidence for and against the null hypothesis. All RT analyses were conducted on the correct responses only (without any outlier exclusion to reflect the full range of variability in the data). We first assessed the results for any age-related potential scaling of the effects, since these have been found not only for overall RT but also for the slowing effect of increased perceptual load, (both are larger for younger children, including for 12 years old as in our sample) in previous selective attention research (Huang-Pollock et al., 2002). Since slower RT can result in scaling of the distractor effects (which was found too in previous research) we first ran a mixed-model ANOVA on the RT in the distractor absent conditions (90% of the trials, to assess these effects unconfounded from any effects of distractor presence) as a function of load and age. This was followed with a mixed model ANOVA on the Distractor Cost on RTs calculated as the proportional increase in RT in the distractor present compared to absent conditions at each load level for each age group. In the sake of completeness, we also report all analyses on the raw RT (i.e. distractor (present/absence) load (low/high) and age group) in the supplementary material.

To provide a measure of sustained attention that corresponds to previous measures of sustained attention in paradigms that did not include distractors, we analysed RT variability in the distractor absent conditions (90% of the trials) as a function of load and age. RT variability was calculated as the coefficient of variation (CV; CV =SDRT/MRT) for distractor absent trials for each participant, in each load level, in order to produce a measure of intra-individual variability which was independent of variations in mean RT and distractor effects.

Due to the count nature of the error data, error rates were analysed using a generalized linear mixed effects model with a Poisson distribution, using the *lme4* package in R. Age group (early adolescence, mid-adolescence, late adolescence and adult), load (low/high) and distractor (absent, present) were included as fixed effects, while participant was included as a random effect. The random effects structure was chosen by comparing the Akaike Information Criterion (AIC) between models that included a random intercept per participant, random intercept for load and participant, or random intercept per participant and random slopes for load (note that models including a random intercept/slope for Distractor effects did not converge). The final model with lowest AIC is reported. The model also included an offset term in order to account for different trial numbers in the Distractor Present (10% of all trials) and Distractor Absent (90% of all trials) conditions. Note that, although the model was fit using error count, the error figure and results section report mean proportions of errors for ease of interpretation in relation to the differences in trial numbers. All post-hoc comparisons were carried out, either correcting the alpha threshold with Bonferroni correction for multiple comparisons for the RT results, or using the *emmeans* package in the error rate analyses, which provides the Bonferroni-Holm corrected p. values. For both RTs and errors the corrections were either made for all contrasts possible, or for the specific contrasts between conditions of interest– these are specified in the results section.

#### Classroom distractibility

Distraction reports were entered into an exploratory factor analysis (EFA) to assess latent distractibility factors. Spearman r was then used for assessing all correlations with the latent distractibility factor since it is less sensitive to outliers. Background variables assessed were age, sex, group work, computer use, generalized cognitive abilities (as measured with the CAT) and interest in the lesson. Distractor effect on RT (% distractor cost) and RT variability as measured with CV (in the distractor absent conditions) in the low load conditions for both measures as a clearer task measure of distractor interference that is free from the effects of perceptual load (which interacted with both measures). Since the distractor effect on the proportion errors did not interact with perceptual load, but the main effect was significant we entered the main effect on errors into the correlations.

## Results

### RT

RT as a function of load and age and distractor conditions are presented in Figure 1.

**Figure 1.**
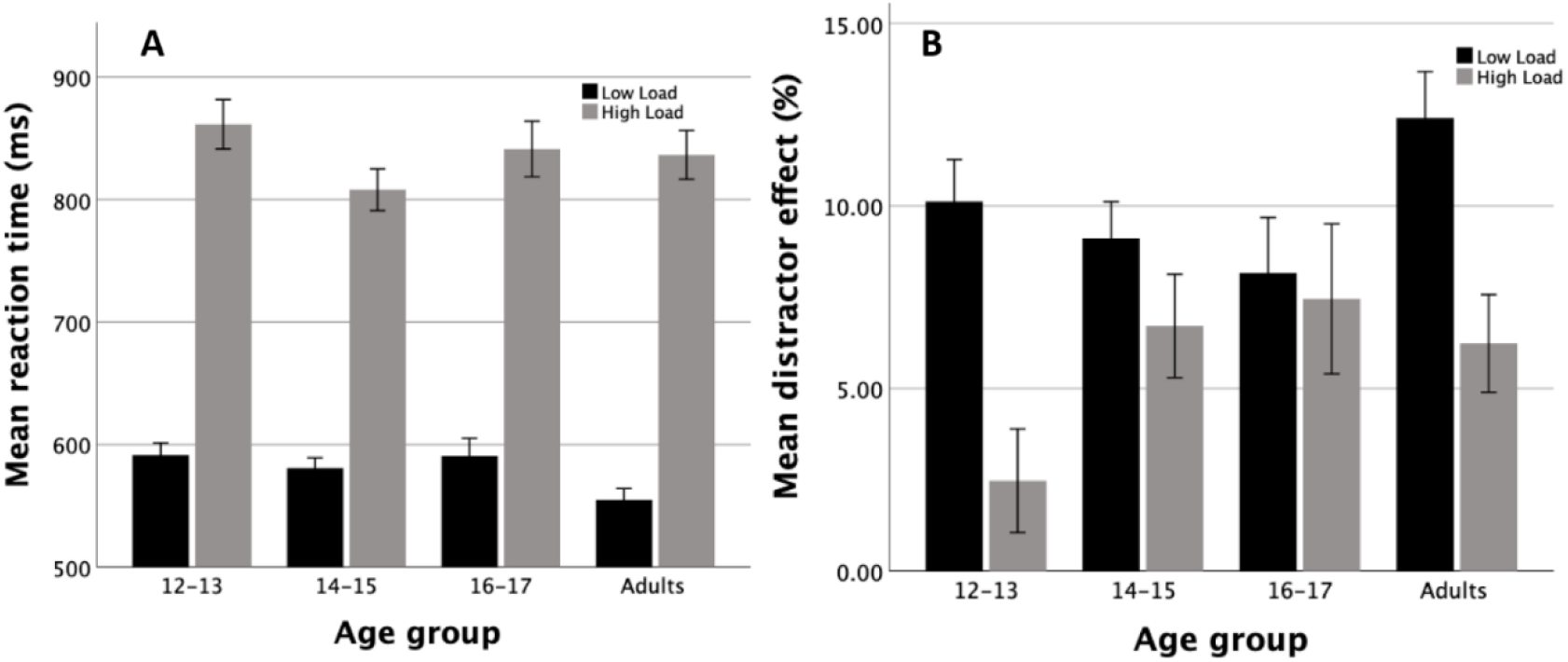
Mean RT as a function of load, age, and distractor conditions. *Note.* Figure 1, panel A shows the mean RT as a function of load and age, in the distractor absent conditions, while the % distractor cost as a function of load and age is shown in panel B. Error bars indicate ±1SE.

To assess whether age led to any effect on the RT or on the RT increase with load (since either of these can lead to scaling of the distractor effect in children, see Huang-Pollock et al., 2002) a mixed-model ANOVA was run on the RT in the distractor absent conditions with load (low/high) as a within-subjects factor, and age group (early adolescence, mid-adolescence, later adolescence, and adults) as a between-subjects factor. The ANOVA revealed a main effect of load, F(1,202) =801.74, p < .001, η^2^_p_ =.799, reflecting longer RT in high load (M = 860, SE =11) compared to low load (M =608, SE =6) conditions. There was no main effect of age (F < 1) however BF_10_ = 0.08 indicated only anecdotal evidence for a null effect. There was an interaction (bordering significance) between load and age group, F(3,202) = 2.61, η^2^_p_ =.04, p = .053, with a BF_10_ of 3.8 indicating substantial evidence for this effect (i.e. for H1). As can be seen in Figure 1A, this interaction reflected a larger load modulation of the RT in the youngest age group compared to the mid adolescents (t(123) = 1.982, p = 0.05, d = 0.36) and no differences between the mid adolescents and late adolescents (t(101) = −1.034, p = 0.304, BF_10_ = 0.36), who also did not differ from the adults (t(79) = −1.293, p = 0.20, BF10 = 0.48, but we note that the BFs for both only indicated anecdotal evidence for a null effect). The greater modulation of load on RT in the early adolescents is as expected from previous research (see Huang Pollock et al. 2002).

To avoid any potential scaling effects of distractor cost due the differing effects of load on RT between the different age groups, we next analysed the distractor cost as the proportional increase in RT in the distractor present compared to absent conditions at each load level (Figure 1B)^3^. A mixed-model 2×4 ANOVA on the proportional distractor RT cost with load (low/high) as a within-subjects factor, and age group as a between-subjects factor, revealed a significant main effect of load, F(1,202) =19.37, p < .001, η^2^_p_ =.09, reflecting a significant reduction of distractor effects found in low load (M =10.0%) under the condition of high perceptual load (M = 5.7%), as expected. There was no main effect of age group, F(3,202) =1.50, p =.22, η^2^_p_ =.02, however BF_10_ = .05 only indicated anecdotal evidence for a null effect, this is likely due to findings that the effect of age group significantly interacted with load, F(3,202) =2.74, p < .05, η^2^_p_ =.04. Figure 2 suggests that this interaction may have been driven in large from the greatest reduction of distractor effects with higher load found in early adolescence, compared to all the older age groups, as expected from previous research as well as a reduction in the load effect in mid to late adolescence groups compared to adults. However none of the post-hoc comparisons of load modulation of distractor effects between the ages passed the Bonferroni-corrected alpha threshold of p=.016 016 (load modulation between early and mid-adolescence groups, t(123)=-2.03, p=.04, d=-.37, BF10 = 1.18, between mid and later adolescence groups, t(101)=-.57, p=.57, d=-.12, BF10 = 0.26, and between later adolescence and adult groups, t(79) =2.09, p =.04, d =.47, BF10 = 1.51). Nevertheless since the increased load modulation of distraction in the early adolescence age compared to the older groups is as expected on the basis of previous research that included 12 years old participants (Huang-Pollock et al., 2002, Study 2), it is more suggestive as an account for the interaction.

**Figure 2.**
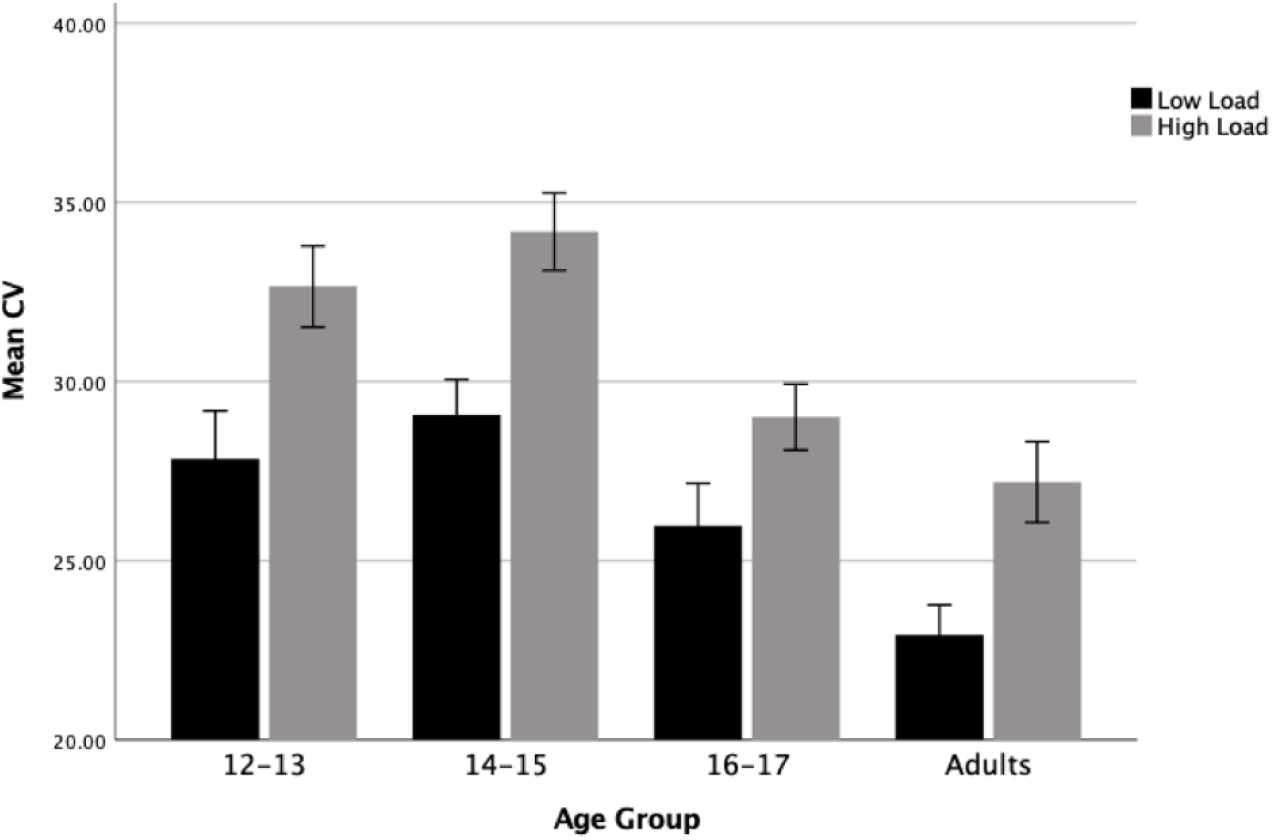
Mean CV of RT as a function of load and age. *Note.* CV as a function of load level and age, for distractor absent trials only as a function of load and age group. Error bars represent ±1SE.

### Response Variability

The results of the CV measure of response variability are shown in Figure 2. To examine sustained attention we ran a mixed-model ANOVA on CVs in the distractor absent conditions with load (low/high) as a within-subject factor and age (early adolescence, mid-adolescence, later adolescence, and adults) as a between-subjects factor. The ANOVA revealed a main effect of load F(1,202) =35.41, p < .001, η^2^p =.15) which reflected larger CV in high load (M =0.31, SE =.6) than in low load (M =0.27, SE =.6) conditions, across all ages. There was a main effect of age, F(3,202) =11.38, p < .001, η^2^p =.15, which did not interact with load (F < 1, although BF_10_ = 0.05 indicating only anecdotal evidence for a null effect). As can be seen in Figure 2, and confirmed with post-hoc comparisons with the Bonferroni corrected alpha of p =.016, there was no significant difference between early and mid-adolescence CV (early adolescence M =0.31, mid-adolescence M=0.32, t(123) =-.47, p =.64, *d* =-.09, BF10 = 0.16 indicating substantial evidence for a null effect), however the late adolescence group had reduced CV (M =0.28) compared to the mid-adolescence group (t(101) =2.74, p =.007, *d* =0.58), and larger CV compared to adults (M =0.25), though this difference did not reach the corrected significance threshold (t(79) =1.98, p =.05, *d* =.45, and BF_10_ =1.39, only suggested anecdotal evidence for this effect).

A 4 (age group) × 2 (load) × 6 (block position) mixed model ANOVA was also run to assess any potential effect of time on task on the CV(see supplementary Figure 3). This analysis showed no significant main effect for block (F(1, 198) = 1.47, p = 0.23, BF_10_ < 0.01, or any interaction with block position (Block × Age Group: F(3, 198) = 0.82, p = 0.48, η^2^_p_ = 0.01, BF_10_ < 0.01, Block × Age Group × Load: F(3, 202) = 0.59, p = 0.62, η^2^_p_ < 0.001, BF_10_ < 0.01, and Block × Load: F(1, 202) = 2.50, p = 0.12, η^2^_p_ = 0.01, BF_10_ = 0.08) with all BFs indicating decisive (or strong in the case of the interaction of load × block) evidence for the null hypothesis

### Errors

Figure 3 displays the mean proportional errors as a function of the experiment conditions. A generalized linear mixed effects model on the error rate, revealed a significant effect of Load (Incidence Rate Ratio (IRR) = 2.71, 95% CI [2.47, 2.97], p < 0.001), indicating an increase in error rates in high load (M= 0.25) compared to low load condition (M = 0.10), as expected. There was also a significant effect of Distractor (IRR = 1.17, 95% CI [1.05, 1.31], p = 0.006), indicating an increase in error rates in the distractor present (M = 0.19) versus distractor absent (M= 0.16) conditions. There was no interaction between Load and Distractor (IRR = 1.02, 95% CI [0.89, 1.17], p = 0.745), or Age Group and Distractor (IRR = 1.15, 95% CI [0.92, 1.43], p = 0.234), and no three-way interaction between Load, Distractor and Age Group (IRR = 1.04, 95% CI [0.80, 1.35], p = 0.796). Age had a significant main effect (IRR = 0.51, 95% CI [0.43, 0.62], p < 0.001), reflecting a linear trend for a decrease in error rates as age increases. Post-hoc contrasts with Bonferroni-Holm corrected p-values for 3 tests clarified that while error rates did not significantly differ between early and mid-adolescents (M = 0.21 an M = 0.20 respectively; z = 0.903, p = 0.367), error rate was significantly higher in mid-adolescents compared to late adolescents (M = 0.20 and M = 0.15 respectively; z = 2.911, p = 0.011), who did not differ significantly from the adults (M = 0.13, z = 1.886, p = 0.119). There was also a significant interaction between Age Group and Load (IRR = 1.25, 95% CI [1.05, 1.50], p = 0.015), indicating a larger effect of load in the younger compared to older age groups, although post-hoc comparisons of the load effect between age groups (corrected with Bonferroni-Holm method for 6 tests) did not reveal any significant effects. The only comparison approaching significance was of the load effect between early adolescents and adults (M = −0.155 and M = −0.136 respectively, z = 2.360, p = 0.1097).

**Figure 3.**
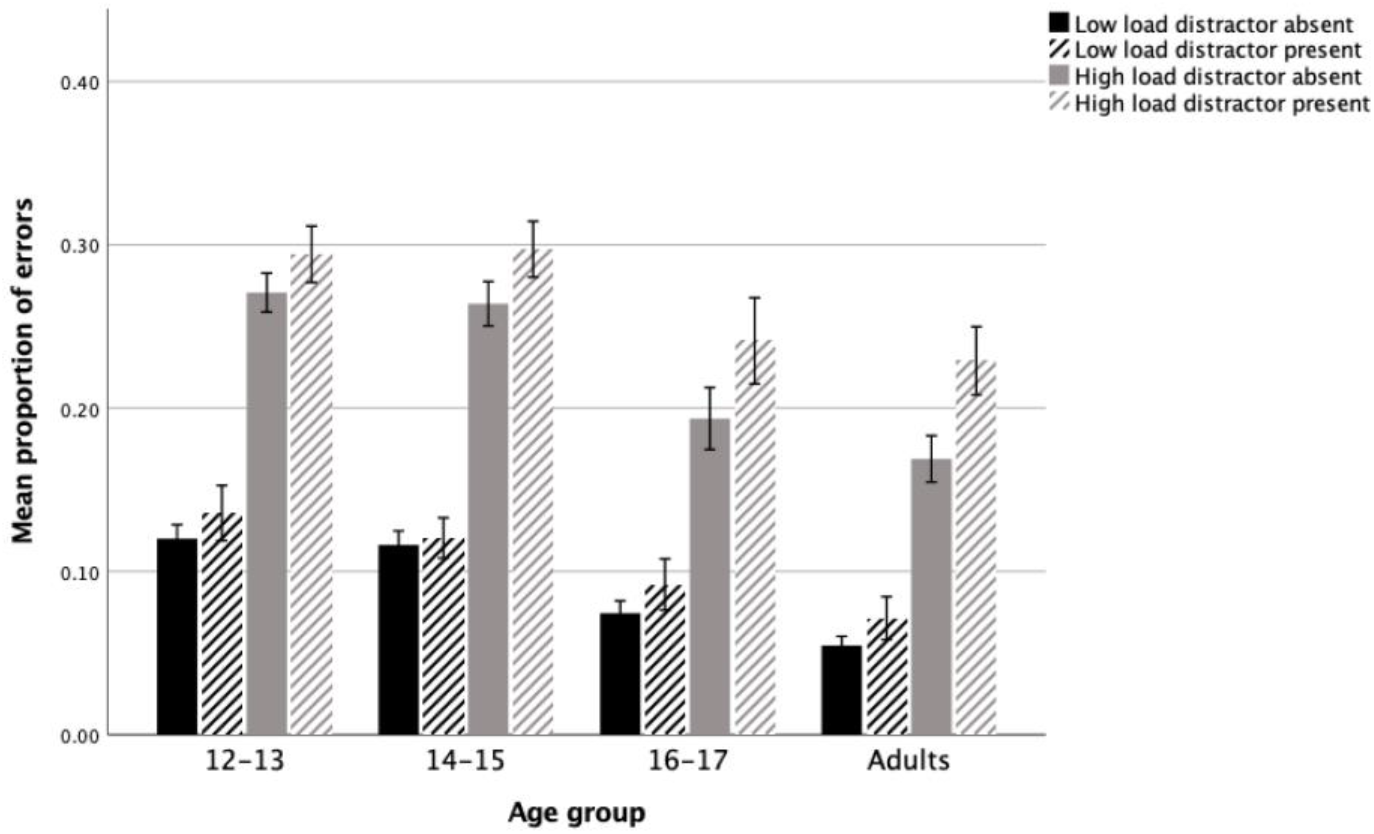
Mean Percentage Accuracy as a function of distractor presence, load and age. *Note.* Mean error proportions given as a function of load level and age. Error bars represent ±1SE.

### Adolescent Distraction During Classroom Learning

Mean distraction duration reports across all distraction sources was 7% (range 0-45%, SD =7%). Exploratory factor analysis of the distraction reports revealed that all distractions loaded onto a single factor at above 0.4, the threshold suggested for the practical significance of a loading value (Stevens, 2009). The scree plot (Figure 4) revealed an inflexion after the first factor which, together with the factor loadings, suggested a single ‘classroom distractibility’ latent variable could be formed for further analysis (see Appendix Table 1 for all the zero-order correlations). This is consistent with previous findings that attention lapses, including both mindwandering and distractibility reported from external distractors, can be described as arising from a single underlying construct when measured in everyday settings (Hobbiss et al., 2019).

**Figure 4.**
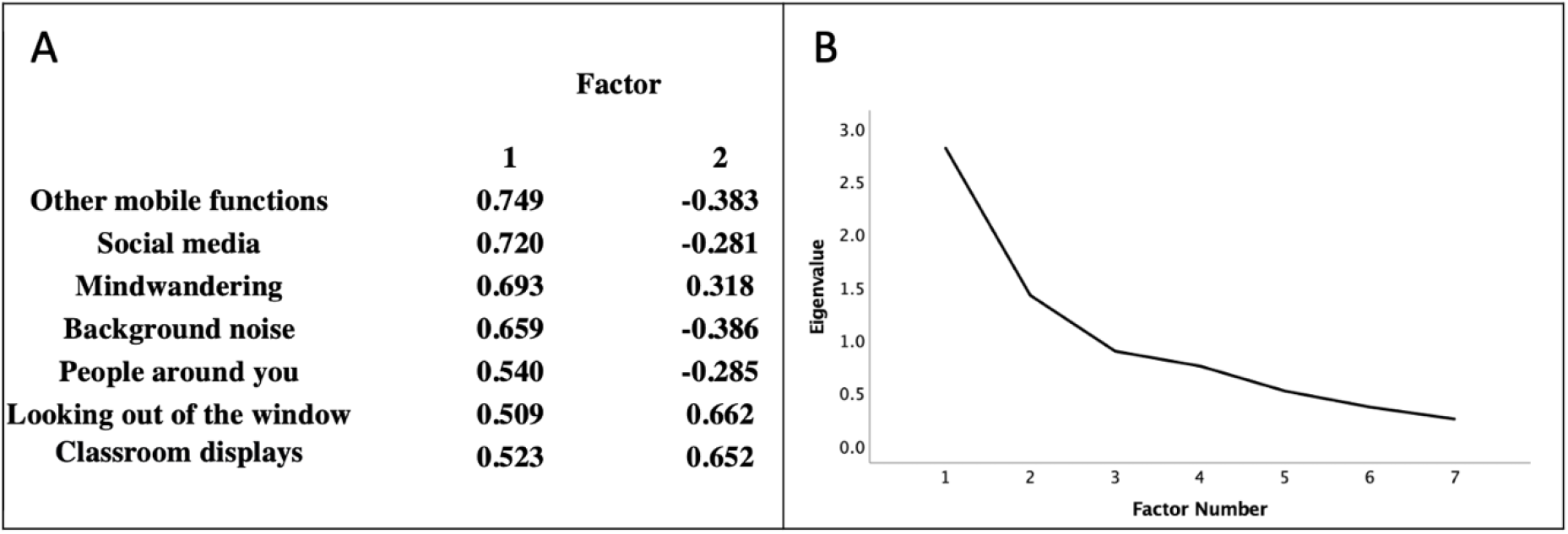
Factor loadings and Scree plot for EFA on distraction variables in adolescents. *Note*. Component loadings (panel A) and scree plot (panel B) for the EFA of the distraction sources reported from the last classroom lesson for adolescent school students. As all distraction reports were positively correlated with one another, an oblique (direct oblimin) rotation was used.

**Table 1.** Multiple regression model for factors predicting classroom distractibility.

| | $\beta$ | T | p | VIF |
| --- | --- | --- | --- | --- |
| **Computer use** | -0.07 | -0.83 | 0.41 | 1.06 |
| **Age group** | 0.04 | 0.43 | 0.67 | 1.15 |
| **Sex** | 0.07 | 0.90 | 0.37 | 1.08 |
| **Group work** | 0.08 | 0.94 | 0.35 | 1.12 |
| **CAT aptitude score** | -0.15 | -1.78 | 0.08 | 1.09 |
| **Interest in lesson** | -0.16 | -2.06* | 0.04 | 1.03 |
| **Distractor RT effect** | 0.18 | 2.16* | 0.03 | 1.16 |
| **Response variability**<br>**(CV)** | 0.26 | 3.29** | 0.001 | 1.06 |
| **Model Statistics** | **R<sup>2</sup>** | **Adjusted R<sup>2</sup>** | **F** | **p** |
|  | 0.173 | 0.124 | 3.53** | <.001 |

*Note.* $\beta$ =Beta, the standardized regression coefficient; Adjusted R<sup>2</sup> significance levels are for R<sup>2</sup> change F-tests

Spearman correlations indicated that distractor interference effects and RT variability in the low load conditions were both significantly correlated with the ‘classroom distractibility’ latent variable: greater distractor interference effect on RT, and greater RT variability in the task performance, were both associated with greater duration of distractibility reported in the classroom (*r_s_* =0.18, p =.03; *r_s_* =.27, p < .01, respectively; See Figure 5). CAT score was significantly negatively correlated with distractibility in the classroom (*r_s_* = −0.19, p =.02). The load modulation effect (the difference between the % distractor RT cost in high load compared to low load) and the main effect of distractor interference on error counts showed no significant correlation with the classroom distractibility latent factor (*r_s_* =.06, p =.5, and *r_s_* =.00, p =.9 respectively) and so were not analysed further.

**Figure 5.**
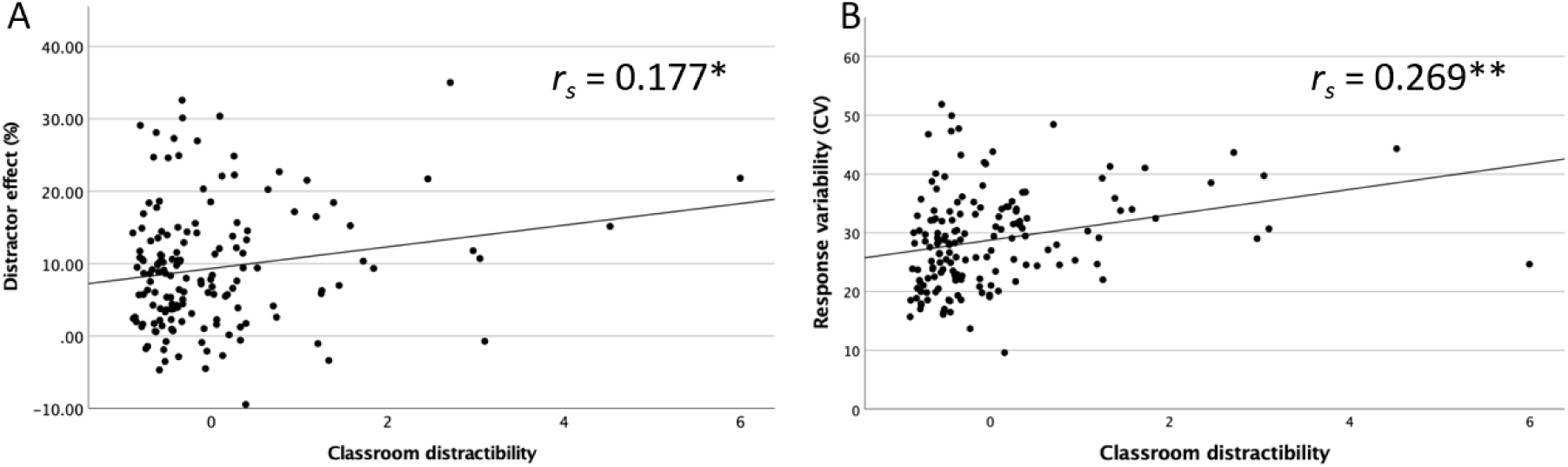
Scatterplots for correlations of experimental variables with classroom distractibility latent variable. *Note.* Classroom distractibility refers to EFA factor scores

A simultaneous regression analysis (Table 1) was conducted to assess whether classroom distractibility latent variable could be predicted from experimental performance, when controlling for background variables (age group, sex, group work, computer use, generalized cognitive abilities and interest in the lesson). Age group and sex were dummy coded.

This analysis revealed that both distractor RT interference and response variability (CV), were significant positive predictors of Classroom Distraction, when controlling for all other variables. Interest in the lesson was also a significant (negative) predictor of classroom distractibility. A further level of the regression model included interaction terms between age and distractor RT effect, CV and interest in the lesson. No interaction terms were significant, so they were omitted from Table 1. The variance explained by CV was largely independent of that explained by distractor interference; distractor interference explained 2.3% and CV 6.3% of the variance in classroom distractibility (above and beyond all control variables) when entered individually, and 7.8% when entered together. As the variance explained by the two together only exceeded the sum of the variance that each factor explained individually, by .08% they represent relatively independent predictors of distraction in our model. In addition, they were also not significantly correlated with one another, *r_s_* =-.11, p =.13. Interest in the lesson was not correlated with either of these variables (*r*_s_ Distractor RT effect =-.02, p =.8; *r*_s_ CV =.04, p =.6), and explained 1.5% of unique variance. It is notable that both experimental variables (Distractor RT effect and CV) appeared therefore to be stronger predictors of classroom distraction than interest in the lesson.

## General Discussion

The present findings established that adolescence involves reduced ability to sustain a stable level of attention focus as reflected in their higher level of response variability. The findings demonstrated that the ability to sustain attention on the task continued to develop throughout adolescence into adulthood. Moreover, this developmental trajectory was found throughout task performance and irrespective of the level of attentional demand in the task (i.e. across both conditions of low and high perceptual load).

In contrast, adolescent’s selective attention ability as measured with the magnitude of distractor interference effects did not generally differ from adults, suggesting selective attention has matured by early adolescence. A more specific developmental trajectory has been found instead, in the effects of perceptual load on distractor processing, which was found to be most pronounced in the early adolescence period. Specifically, increased perceptual load led to a larger reduction in distractor processing in early adolescents (and to a small extent also mid adolescents) compared to older adolescents. A larger impact of perceptual load on task RT (even in the absence of a distractor) and error rates was also found in younger adolescents compared to older adolescence. This pattern of findings suggests that while selective attention per se (i.e. the general ability to ignore irrelevant distractors) has matured by the adolescence period, perceptual capacity that is known to develop in younger children (e.g. Huang-Pollock et al., 2002) continues to develop in the early to mid-adolescence period.

Importantly, both measures of sustained attention and selective attention were established as significant unique predictors of adolescents self-reported experience of distractibility during a classroom lesson, thus establishing their external validity, and applicability to prediction of attention focus in the important educational settings of the school classroom. We discuss all these findings in turn below.

### Selective Attention in Adolescence

The findings that development of selective attention specifically involved the effects of perceptual load on the ability to focus attention in the younger adolescence period rather than reduced ability to ignore irrelevant distractors, is generally consistent with previous research, and clarifies the impact of perceptual load on selective attention during adolescence, as follows.

The findings of a generally comparable magnitude of distractor interference effects across adolescents of all ages and adults is consistent with previous research reports of a similar ability to ignore distractors between adolescents and adults in studies using the response competition flanker task paradigm (Grose-Fifer et al., 2013; Gyurkovics et al., 2020; Huizinga et al., 2006). The present study extends the support for the conclusion that selective attention per se has matured by the adolescence period to the case of salient and meaningful yet entirely irrelevant distractor faces.

The contrast between the lack of developmental effects with these types of distractors and the previous findings of reduced focused attention in adolescence when affective or motivationally-relevant distractors are presented (e.g. Grose-Fifer et al., 2013; Roper et al., 2014) suggests that adolescence involves an interactive impact of the development of control over distraction by motivationally-relevant or more affective stimuli, rather than reduced ability to focus attention per se, in the presence of any distractors. This is however a speculative suggestion that awaits further testing, contrasting development of selective attention within an irrelevant distractor task in which nature of distractors is manipulated, comparing affective or reward-associated distractors with similarly salient distractors that are both emotionally neutral and have not been associated with a reward. Such a paradigm would also have the advantage of testing for an interaction of age and distractor type, rather than basing a conclusion on a null finding (lack of developmental effect).

Our suggestion that selective attention has matured by early adolescence together with our findings that increased perceptual load in our task reduced distractor processing (as long as the scaling of RT by increased perceptual load and its modulation by age is taken into account)^4^ are also generally consistent with findings (Couperus, 2011) that increased perceptual load in a task reduces the P100 evoked potential related to distractor processing to an equal extent across children (aged 7-9, 10-12), adolescents (of 13-15, and 16-18 age groups) and adults (over 19 years old). Since the P100 is considered a sensory component these findings suggest that the early-selection mechanism that allows for perceptual filtering through reduced sensory-cortex responses is fully mature already in late childhood and early adolescence. Importantly, Load Theory makes a clear dissociation between early (perceptual) selection attention mechanism that benefits from increased perceptual load, and late-selection cognitive control mechanisms (associated with frontal cortex ‘executive control’ regions) such as working memory that serves to maintain performance in line with the task-priorities (e.g. Lavie et al., 2004). This dissociation can further clarify the contrasting findings of immature selective attention in tasks involving affective or motivation-related distractors, by attributing these to the continued development of the late-selection mechanisms (associated with the continued maturation of the frontal cortex cognitive control regions). Since this type of distractors are stronger competitors for attention than the target, their suppression places greater demand on the immature cognitive control mechanisms.

Finally, our results of a developmental trajectory in the effects of perceptual load on selective attention suggest that while early-selection filtering may be fully developed in early adolescence, perceptual capacity is still smaller in the younger age groups (especially our early adolescents group) who displayed greater load modulation of both RT and distractor effects on RT. This is consistent with previous findings within the flanker task paradigm (Huang-Pollock et al., 2002) and suggests that perceptual capacity continues to develop in early adolescence (see also Remington et al., 2014, for a demonstration that perceptual capacity may also continue to develop beyond early adolescence). Specifically, Huang-Pollock et al. (2002) demonstrate that a smaller capacity is exhausted already by smaller increases in perceptual load (as predicted from the resource approach underlying load theory). The greater reduction of distraction in younger adolescents by the same level of high perceptual load found in our study can be similarly explained by the high task leaving less ‘spare’ resources to spill over to distractor processing in younger adolescents because of their smaller capacity (compared to older adolescents).

Couperus (2011) also found a larger load modulation in younger versus older children with the younger children requiring longer presentation time of targets and this corresponds to our findings of a greater load modulation of the RT. Although Couperus (2011) did not find an interaction of age and load with their selective attention measure (both behavioural and ERP) this may be due to reduced sensitivity of the selective attention measure used in the study (where the unattended probe did not appear at the same time as the target) compared to the distractor measures used here and in Huang-Pollock et al. (2002) study.

### Selective attention and inattention in the classroom

The magnitude of distractor interference effects on task performance was found to predict the duration of reported distractibility in the school classroom. Moreover, distractor interference was a significant predictor when controlling for age, gender, interest in the topic of study, computer use, and group work, as well as when controlling for generalized cognitive abilities (as measured with CAT score) in adolescence. These findings demonstrate that the measure of focused attention through the interference effects produced by task-irrelevant distractors clearly does not depend on immature development for its prediction of classroom distraction across all ages tested.

In addition, the findings that adolescents’ susceptibility to distraction both from a variety of external sources (e.g. background noise, mobile devices), and from internal sources in the form of mindwandering, all converged on a latent factor of classroom distractibility which could be predicted from our task measure of distractor interference suggests that our measure reflects a fundamental trait-like mechanism of focused attention. Previous research has established that the magnitude of distractor interference effects in adults correlates with their reports of mindwandering in daily life (Forster & Lavie, 2014), and adult self-reports of childhood inattentive symptoms (Forster & Lavie, 2016). The present results extend this pattern to the important situation of adolescents’ experience of distractibility during their school lessons.

The extension to the school classroom also complements previous findings that behavioural task measures (such as Stroop-like paradigms, requiring response inhibition) can correlate with teacher ratings of students’ attention control skills (at least in terms of differentiating between the top and bottom quartiles for children up to age 12 (Das et al., 1992; Papadopoulos et al., 2002). Importantly, these findings advance previous research by clarifying that the prediction of attention in the classroom cannot be explained any of the factors controlled for in our regression model, at least when the measure is based on the pupils’, rather than teachers’, report. An important factor is the CAT scores. These were found to correlate with attention focus in the classroom, but not significantly when attention measures are included in the simultaneous regression model, while the distractor interference remained a significant predictor of classroom inattention when CAT scores were controlled for. Considering also the well-established correlations of intelligence and cognitive abilities measures and educational achievement (see Roth et al., 2015 for a meta-analysis), one plausible account for our pattern of findings may be that the ability to focus attention is a critical factor both for education in the classroom and for performance of cognitive abilities and intelligence tests. Although rather speculative at present, this suggestion is consistent with previous findings that sustained attention tasks (error rates in a task which required the continuous solving of easy arithmetic problems) predicts academic performance above and beyond intelligence (Steinmayr et al., 2010).

### Sustained Attention in Adolescence

Adolescence was found to involve a clear developmental trajectory in the ability to sustain stable levels of attention, as measured with response variability in the attention task in the distractor-absent conditions. Adolescents of 12-15 years of age showed greater response variability compared to 16-17 year olds, who in turn had a trend for greater response variability compared to adults.

This developmental pattern was found throughout task performance, with BFs indicating decisive (or strong in the case of the interaction of load x block) level of evidence for a lack of effect of time on task. This suggests adolescents have immature ability to sustain stable levels of attention during task performance, rather than a specifically reduced ability to sustain attention as the time goes on in the task. Although we note that the lack of effects of time on task on adult performance too might simply suggest this task was not sensitive to reveal any extra component of time on task effects on sustained attention. Future research with longer task version in our paradigm may prove useful in assessing the effects of time on task. Importantly the developmental effects on sustained attention were found across perceptual load levels, even though high perceptual load was shown to be effective in increasing the ability to focus selective attention, as shown by reduced distractor interference effects for adolescents of all ages. These findings are consistent with previous research showing increased response variability in adolescents’ performance of sustained attention tasks such as the SART or CPT (Braet et al., 2009; Stawarczyk et al., 2014; Fortenbaugh et al., 2015; Gyurkovics et al., 2020) and extend it to establish that the ability to sustain attention is still developing in adolescence, even in tasks of high perceptual load that clearly improve their ability to focus selective attention and ignore irrelevant distractors similarly to adults.

The contrast we establish between selective and sustained attention, when both are measured within the same task allows us to rule out alternative accounts for our findings of immature sustained attention in adolescence, in terms of any general non-attentional factors (e.g. reduced motivation to pay attention to the task), since these should result in similar effects on our two attention measures. For example, an account of the reduced ability to sustain attention on the task attributing it to immature ability to sustain the same level of motivation throughout performance of rather monotonous tasks (such as CPT and SART), should also predict reduced selective attention, manifested as increased distractor processing in the adolescent participants. This is especially so given that our irrelevant face distractors carry more interesting socio-biological information than the task letters, and thus motivational factors should have had a more pronounced effect on adolescence levels of motivation to ignore these more interesting distractors, but this was not the case.

In addition, any accounts of the increased variability in our task in terms of generalised immaturity of control over the response execution or temporal coordination of responses specifically leading to the variability in RTs (which may apply to effects seen in tasks involving congruency sequential effects, (CSE, e.g. Gyurkovics & Levita, 2021) are unlikely to provide a compelling alternative account for the increased response variability in adolescence found in our task, for the following reasons. Our task did not involve varying levels of response conflict, which drive RT variability in their own right in relation to the specific trial sequences. Moreover, our CV measure was based on the majority of the task trials (90%) which involved routine choice response to one target letter or the other, and thus mostly unaffected by any slowing effects due to the minority of trials in which there was a distractor. Finally, and most importantly, our CV measure was found to significantly correlate with adolescents reports of inattention in the classroom, including specially also a strong correlation with mindwandering reports. The correlations with classroom inattention provide strong support for our proposal that increased CV in our task reflects reduced ability to sustain stable levels of attention on the task as reported also in previous sustained attention research (e.g. Cheyne et al. 2009) and as we discuss further next.

#### Sustained attention and inattention in the classroom

The present results also establish our sustained attention measure as a predictor of adolescents’ ability to focus attention in the classroom, including when other factors (such as age, gender, computer use, group work, CAT, and interest in the lesson) are controlled for. Our findings that response variability predicts inattention in the classroom when interest in the lesson is controlled for (in itself also a predictor of inattention, as expected) further supports the conclusion that our measure reflects immature attention rather than reduced motivation to engage in tasks of little interest. These findings are consistent with earlier reports (Cheyne et al. 2009; Stawarczyk et al., 2014) that response variability during SART correlates with adolescents reports of both mindwandering, and with reported cases of attention failures in daily life. Although Stawarczyk and colleagues did not find a correlation between response variability and external distractions this can explained with the reduced sensitivity to assess external sources of distraction in the rather austere conditions of the laboratory.

There are a number of potential causes for the relationship found here in adolescents between increased variability and greater distractibility during learning. A compelling account may be made on the basis of the well-known protracted development of executive control and the associated frontal control networks in adolescents. Fluctuations of executive control may lead to both increased response variability on the task and vulnerability to distraction in the classroom. Indeed ‘task-positive’ prefrontal cortex networks associated with attention control in line with the task demands undergo prolonged maturation during adolescence (e.g., Choudhury et al., 2008; Constantinidis & Luna, 2019). These networks have also been found to be important for sustaining attention on task: greater levels of response variability are associated with reduced regulation of spontaneous neural fluctuations by the task-positive networks (e.g., Esterman et al., 2014) as well as being negatively correlated task-related signals in frontal and parietal cortices. The reduced task-positive regulation in frontoparietal networks can explain not only the increased response variability but also the greater vulnerability to distraction in the classroom, attributing both to fluctuations of cognitive control. On this account one might expect to find an effect of age on the distractor interference effects on task, however given that our distractors were less motivationally-relevant than the potential sources of distraction in the classroom, it is plausible that immaturity of cognitive control specifically concerns distractions by motivationally-relevant stimuli, which require greater exercise of cognitive control in order to be ignored.

Attributing sustained attention to immature cognitive control is in accord also with the proposed dissociation offered by Load Theory (e.g. Lavie et al., 2004) between early selection attention which improves by high perceptual load and cognitive control which is shown here to continue to develop. The unique sources of variance explained by each of the sustained and selective attention task measures is in further support of this dissociation.

### Errors

Younger adolescents (age 12-15) were found to differ from older adolescents and adults in producing higher error rates, while the older adolescents performed at adult levels. The findings of improved accuracy with age is generally consistent with previous research, (e.g. Aïte et al., 2018) though different tasks and measures differ somewhat with respect to whether older adolescents reach adult levels of performance accuracy already at age 16-17 (e.g. Leon-Carrion et al., 2004; or even somewhat earlier e.g. at age 15 in Huizinga et al., 2006); or continue to improve up to age 21-22 (e.g. Duell et al., 2018; Stawarczyk et al., 2014).

It is plausible that performance accuracy in the latter tasks is more sensitive to reveal any remaining developmental trajectory in older adolescents, compared to our task, either due to being more monotonous (e.g. as in the SART) and thus tapping more on immature sustained attention abilities, or involving greater demand on response inhibition (due to their involvement of no-go targets).

## Conclusions

The present research establishes that selective attention matures earlier in adolescence compared to sustained attention. The main developmental trajectory found for selective attention was a greater modulation of both task RT and distractor interference by increased perceptual load in the early adolescence period potentially indicating continued maturation of perceptual capacity previously established in earlier childhood. However no difference in the general magnitude of distractor interference effects were found across the sample, suggesting early selection distractor filtering mechanism have matured by adolescence. In contrast our finding showed that adolescents’ ability to sustain attention is still developing up to and including age 16-17, as indicated by elevated levels of response variability. Importantly our research established the developmental dissociation between mature selective attention and continued development of sustained attention within the same task and under conditions of high perceptual load that improved their attention focus to the same extent as in adults. Moreover both our task measures of selective attention and sustained attention were found to be unique predictors of adolescents’ inattention levels in the school classroom, even when important background factors including cognitive aptitude and level of interest in the lesson are controlled for, thus establishing the external validity of our attention task measures and their potential applicability to the understanding of attention in the important educational settings of the school classroom.

Since each of the sustained and selective measures predicted a unique source of variance in classroom inattention, and were dissociable also in their development our results suggest that sustained attention may depend on the development of cognitive control and the associated frontal cortex maturation, while selective attention and focusing on tasks under more attention demanding conditions (of high perceptual load) depends on the ‘early selection’ mechanisms of focused attention (Lavie et al., 2004), and these have matured by the adolescence period, although the younger adolescents still have a smaller perceptual capacity.

## Author Note

All data is available at the Open Science Framework (OSF) at https://osf.io/cbd4x/

We have no conflict of interest to disclose. We thank Tom Phillips and Ursule Taujanskaite for help with the data collection and analyses, Jenni Kemp for liaising with teachers, and Damian Haigh, Amy Jones, Adrian Cook and the student volunteers for their assistance in our host school.

---

**Appendix Table 1.**
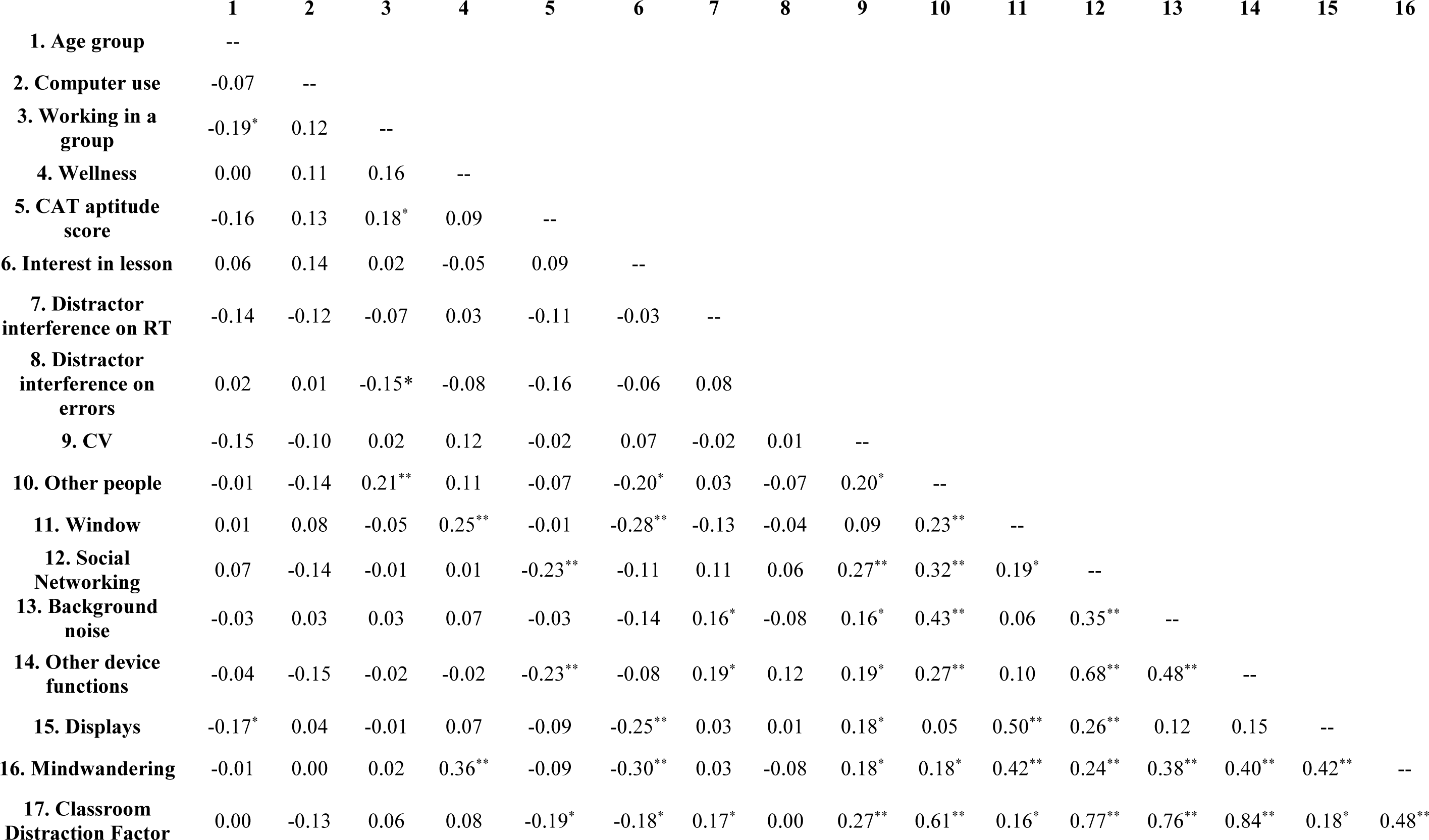
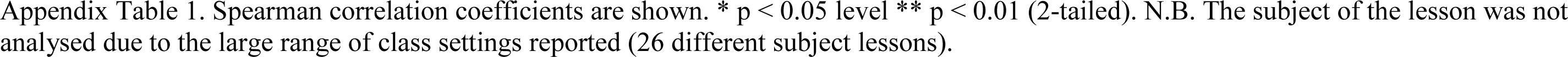
Zero-order correlations between all variables of interest – Adolescents.

Spearman correlation coefficients are shown. \* $p < 0.05$ level \*\* $p < 0.01$ (2-tailed). N.B. The subject of the lesson was not analysed due to the large range of class settings reported (26 different subject lessons).
|  | 1 | 2 | 3 | 4 | 5 | 6 | 7 | 8 | 9 | 10 | 11 | 12 | 13 | 14 | 15 | 16 |
| --- | --- | --- | --- | --- | --- | --- | --- | --- | --- | --- | --- | --- | --- | --- | --- | --- |
| **1. Age group** | -- |  |  |  |  |  |  |  |  |  |  |  |  |  |  |  |
| **2. Computer use** | -0.07 | -- |  |  |  |  |  |  |  |  |  |  |  |  |  |  |
| **3. Working in a group** | -0.19* | 0.12 | -- |  |  |  |  |  |  |  |  |  |  |  |  |  |
| **4. Wellness** | 0.00 | 0.11 | 0.16 | -- |  |  |  |  |  |  |  |  |  |  |  |  |
| **5. CAT aptitude score** | -0.16 | 0.13 | 0.18* | 0.09 | -- |  |  |  |  |  |  |  |  |  |  |  |
| **6. Interest in lesson** | 0.06 | 0.14 | 0.02 | -0.05 | 0.09 | -- |  |  |  |  |  |  |  |  |  |  |
| **7. Distractor interference on RT** | -0.14 | -0.12 | -0.07 | 0.03 | -0.11 | -0.03 | -- |  |  |  |  |  |  |  |  |  |
| **8. Distractor interference on errors** | 0.02 | 0.01 | -0.15* | -0.08 | -0.16 | -0.06 | 0.08 | -- |  |  |  |  |  |  |  |  |
| **9. CV** | -0.15 | -0.10 | 0.02 | 0.12 | -0.02 | 0.07 | -0.02 | 0.01 | -- |  |  |  |  |  |  |  |
| **10. Other people** | -0.01 | -0.14 | 0.21** | 0.11 | -0.07 | -0.20* | 0.03 | -0.07 | 0.20* | -- |  |  |  |  |  |  |
| **11. Window** | 0.01 | 0.08 | -0.05 | 0.25** | -0.01 | -0.28** | -0.13 | -0.04 | 0.09 | 0.23** | -- |  |  |  |  |  |
| **12. Social Networking** | 0.07 | -0.14 | -0.01 | 0.01 | -0.23** | -0.11 | 0.11 | 0.06 | 0.27** | 0.32** | 0.19* | -- |  |  |  |  |
| **13. Background noise** | -0.03 | 0.03 | 0.03 | 0.07 | -0.03 | -0.14 | 0.16* | -0.08 | 0.16* | 0.43** | 0.06 | 0.35** | -- |  |  |  |
| **14. Other device functions** | -0.04 | -0.15 | -0.02 | -0.02 | -0.23** | -0.08 | 0.19* | 0.12 | 0.19* | 0.27** | 0.10 | 0.68** | 0.48** | -- |  |  |
| **15. Displays** | -0.17* | 0.04 | -0.01 | 0.07 | -0.09 | -0.25** | 0.03 | 0.01 | 0.18* | 0.05 | 0.50** | 0.26** | 0.12 | 0.15 | -- |  |
| **16. Mindwandering** | -0.01 | 0.00 | 0.02 | 0.36** | -0.09 | -0.30** | 0.03 | -0.08 | 0.18* | 0.18* | 0.42** | 0.24** | 0.38** | 0.40** | 0.42** | -- |
| **17. Classroom Distraction Factor** | 0.00 | -0.13 | 0.06 | 0.08 | -0.19* | -0.18* | 0.17* | 0.00 | 0.27** | 0.61** | 0.16* | 0.77** | 0.76** | 0.84** | 0.18* | 0.48** |

## Footnotes

1 We note that the Stroop task has also been used to study attention in adolescence. However, the integration of the target and distractor within the same stimulus in the typical Stroop Colour-Word stimuli precludes any clear interpretation in terms of selective attention, since space-based or object-based separation of target and distractors are known as pre-requisite conditions for successful selective attention (e.g. Lavie & Driver, 1996). Instead Stroop effects can be well accounted for by reduced control over response inhibition which continues to develop during the adolescence period (e.g. Aïte et al., 2016).

2 We administered also a similar distraction checklist to the adult samples (adapted to university lessons), however initial inspection of the student responses revealed that despite our scheduling the experiment following lecture slots, the students did not arrive straight from a lecture (a substantial number of students reported having arrived from home or a computer room). We therefore did not continue with these analyses.

3 The results for distractor effects on the mean raw RTs, as a function of load and age are reported in Supplementary material.

4 We provide the results and related discussion of the findings when scaling of RT (by load as well as the interactive effect of age and load) is not taken into account in the supplementary material.

