## Supplementary material for "Sustained Selective Attention in Adolescence: Cognitive Development and Predictors of Distractibility at School": supp material

**Mean RT analyses**

Mean RT as a function of load and age and distractor conditions are presented in Figure 1.

**Figure 1.**
*Mean RT as a function of load, age, and distractor conditions.*

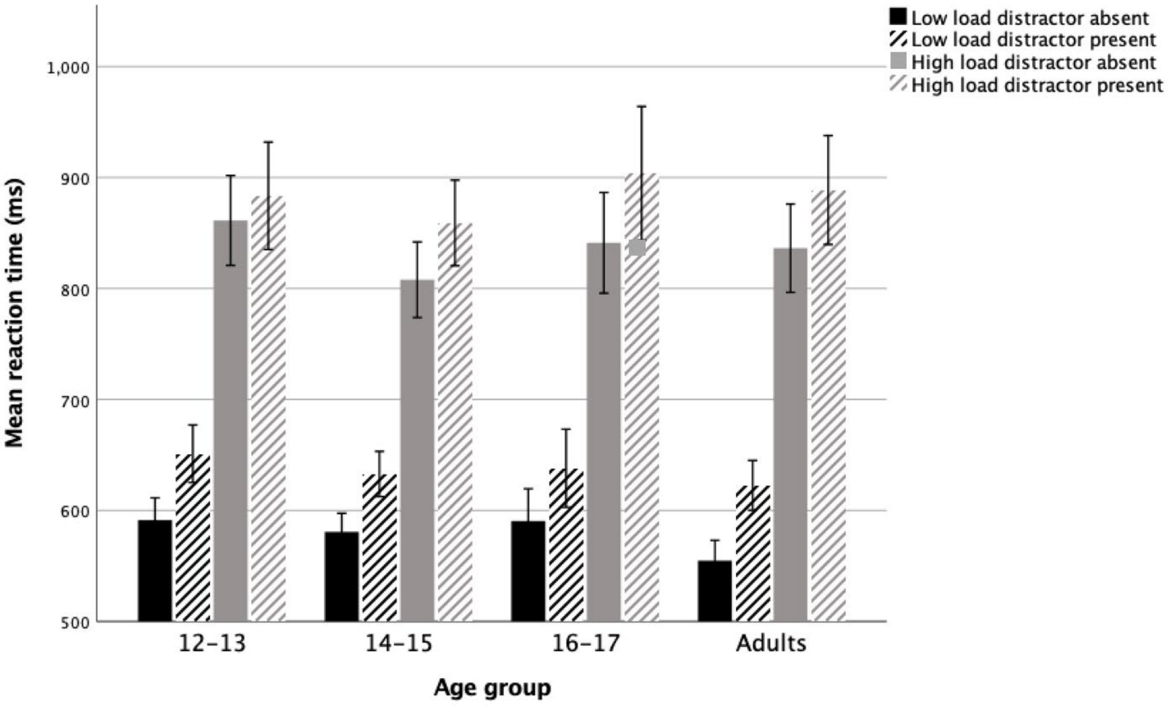


Note. Error bars indicate ±1SE

A 2×2×4 mixed-model ANOVA with load (low/high) and distractor (present/absence) as within-subjects factors, and age group as a between-subjects factor revealed main effects of load, $F(1,202) = 801.74$, $p < .001$, $\eta^2_p = .80$, with slower reaction times in high load (M = 860ms) than in low load (M = 608 ms), and distractor presence, $F(1,202) = 167.97$, $p < .001$, $\eta^2_p = .45$, with slower reaction times in distractor present (M = 760 ms) than distractor absent (M = 708 ms) conditions. There was no main effect of age group, $F < 1$ and $BF_{10} = 0.08$ indicated very strong evidence for a lack of Age effect on the RT. There was no interaction between distractor presence and age group $F(1,202) = 1.08$, $p = .36$, η2p =.02, and $BF_{10} = 0.004$ indicated decisive evidence for a null effect. The interaction between load and distractor presence, $F(1,202) = 1.77$, $p = .19$, $\eta^2_p = .01$. 0.349, was not significant either, however the $BF_{10}$ of 0.35 indicated only anecdotal evidence for a null effect. Although there was also no significant interaction between load and age group, $F(1,202) = 1.51$, $p = .21$, $\eta^2_p = .02$, BF = 3.806 indicated substantial evidence for the H1 hypothesis that there was an interaction. Finally, the three way interaction between load, distractor conditions and age was also not significant, $F(1,202) = 2.23$, $p = .09$, η2p =.03, and BF=.001 indicated decisive evidence for the lack of this interaction and the model comparison analysis showed that the top 10 models did not include this three way interaction. The top 3 models in the BF analyses included the main effects of load and distractor as the top model, as well as the interactions of load by age and load by distractor as the top second and third models respectively.

Overall these results are consistent with the analyses reported in the main paper except for the anecdotal evidence for the lack of load by distractor interaction, and the decisive evidence for a lack of a three-way age x load x distractor interaction since in the percentage distractor cost analyses the analogues two-way interaction of age and load was significant. It is unclear whether these discrepancies are due to the scaling component of the effect of load on RT especially across ages, or whether the presence of distractor faces at fixation is generally subject to reduced modulation by perceptual load and even more so for mid to late adolescents (since Figure 1 shows a clear effect of perceptual load on distractor interference in the early adolescents group) whose attention may have been captured by distractor faces irrespective of increased perceptual load in the task due to their higher sociobiological significance.

Indeed, previous research of attention in adults, established a load modulation of distractor faces only when these are entirely task-irrelevant as here, and presented in the display periphery (Forster & Lavie, 2021). An earlier study (Beck & Lavie 2005) manipulating perceptual load in a response competition paradigm has demonstrated greater processing of distractors when these are presented at fixation, compared to the periphery, and under some conditions weaker (though statistically significant) effect of perceptual load on fixated distractors (Beck & Lavie 2005, Experiment 2). It is therefore plausible that distractor faces at fixation remain stronger competitors for attention than the task stimuli even under conditions of high perceptual load and that adolescents have reduced control over such competition. Future research comparing irrelevant distractor faces in the periphery versus at fixation and other categories of meaningful non-face distractor objects (e.g., Lavie et al., 2003) as a function of age and load is needed to reach a firm conclusion. Importantly for the present study, the distractor effect on the raw RT in the low load conditions was also significantly correlated with the factor of classroom distraction ($r_s$ =.173, p =.03) and was also predictive of the classroom distraction in the simultaneous regression (β = 0.26, t = 2.54, p = .01, VIF = 1.66)

**Figure 2**

*Scatterplot between distractor effect on reaction time and classroom distractibility factor score.*

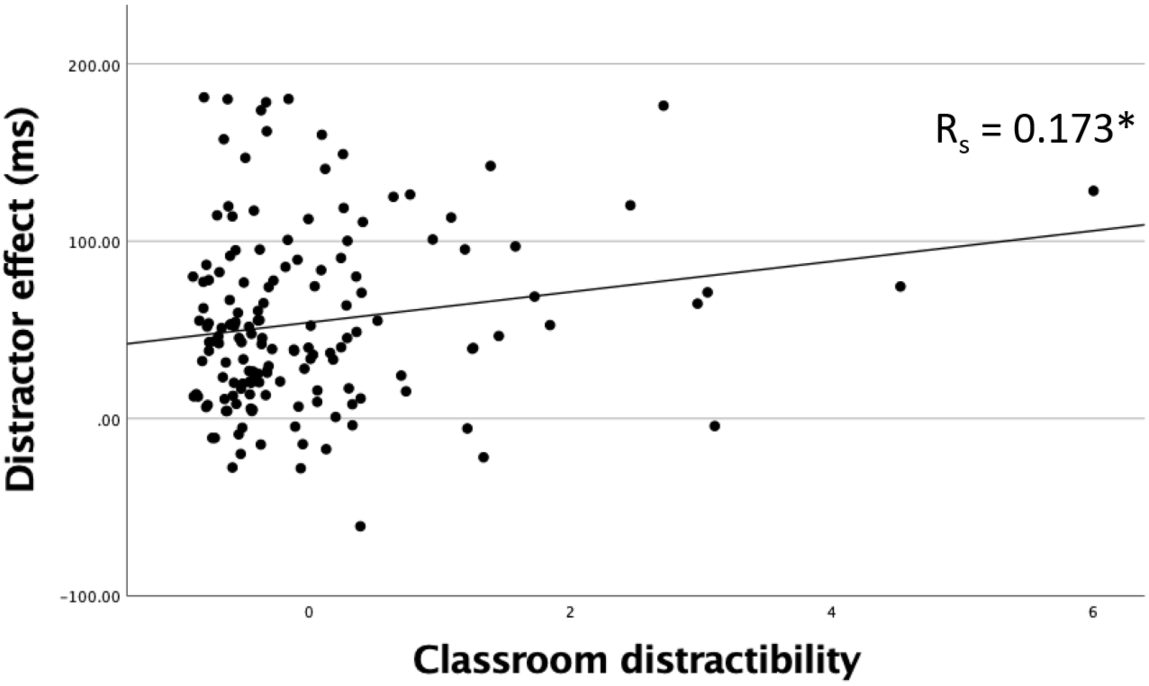


Note. Classroom distractibility refers to EFA factor scores

### CV as a function of time on task

**Figure 3**

*CV in the Attention task as a function of age group, block and load*

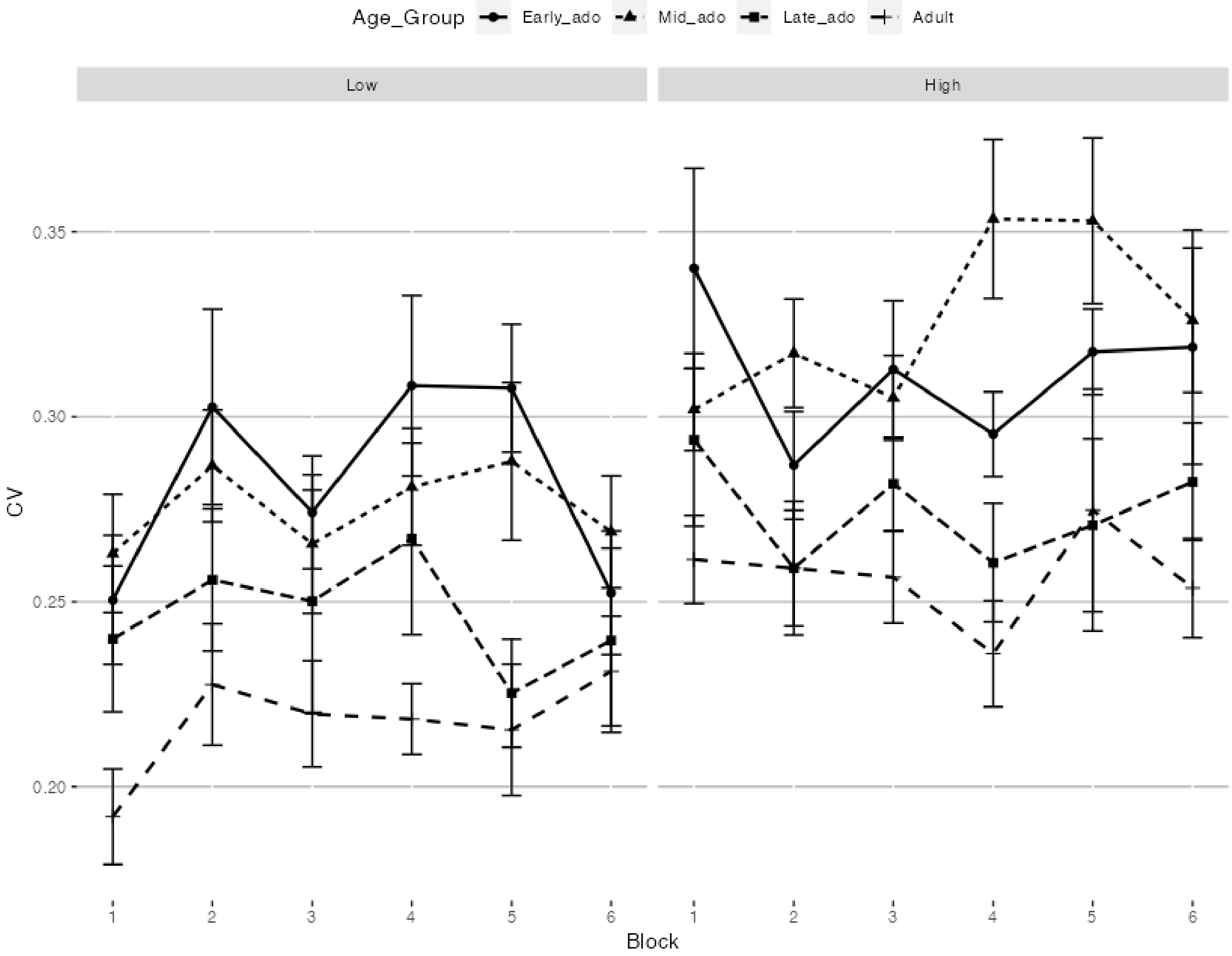


*Note.* Error bars represent ±1SE.
